# Integration of ATAC-seq and RNA-seq identifies novel candidate genes associated with drought tolerance in *Zoysia japonica* Steud

**DOI:** 10.1101/2023.11.26.568709

**Authors:** Liangying Shen, Zewen Qi, Shuwen Li, Mengdi Wang, Jiabao Chen, Jiahang Zhang, Lixin Xu, Liebao Han, Yuehui Chao

**Author notes:** These authors contributed equally (Liangying Shen, Zewen Qi).

## Abstract

**Highlight:** Integration of ATAC-seq and RNA-seq identifies novel candidate genes associated with drought tolerance in *Zoysia japonica* Steud.

The warm-season turfgrass Zoysia japonica is renowned for its drought resistance and serves as an exceptional domestic turfgrass in China. In order to unlock the potential of this native grass, identify drought-resistant genes, enhance the genetic transformation system, and maximize its utilization benefits, we conducted physiological characterization, multi-omics analysis, and RT-qPCR experimental verification in Zoysia japonica. This study suggested that 63 high-confidence genes related to drought stress and 6 motifs regulating drought responses were identified using a combined omics approach and RT-qPCR validation. The study discovered a positive correlation between ATAC-Seq peak intensity and gene expression levels. The expression of high-confidence genes was linked to *Zoysia japonica* resistance evaluation and phenotypic traits, implying that these genes are involved in responding to external drought stress. This study combined ATAC-seq and RNA-seq technologies for the first time to identify drought-related genes in *Zoysia japonica*, elucidating the grass’s adaptation to environmental stress and the regulatory mechanisms underlying stress responses, and laying the groundwork for *Zoysia japonica* improvement and breeding.

## Introduction

Due to global warming, drought stress can harm plant growth and development (Huang *et al*., 2023). Studying the mechanisms underlying drought tolerance in resilient plant species is crucial for addressing the global drought problem and enhancing the drought tolerance of less resilient plants. Plants activate a series of protective mechanisms in response to drought stress in order to mitigate the harm caused by the stress (Wang *et al*., 2022*c*). Turfgrass, for example, increases the activity of protective enzymes in response to drought stress to improve its ability to remove reactive oxygen species (Xu *et al*., 2016; Tan *et al*., 2022). Plants may also actively accumulate osmotic-regulating substances to reduce water loss from cells and improve drought resistance (Ma *et al*., 2021). Drought stress also affects plant metabolic pathways (Xu and Huang, 2018). It includes decreased photosynthesis and respiration, increased organic acid and proline accumulation, decreased chlorophyll content and leaf area, and increased anthocyanin synthesis (Merewitz *et al*., 2011; Ma *et al*., 2018). Furthermore, drought stress has a wide-ranging effect on plant gene expression (Bushman *et al*., 2021; Xu *et al*., 2013). It increases the levels of antioxidant, dehydrin, and proline-synthesizing enzymes while decreasing the levels of genes involved in photosynthesis and chlorophyll synthesis (Merewitz *et al*., 2011; Guan *et al*., 2022*a*). In addition, drought stress can influence gene expression patterns and non-coding RNA expression (Gelaw and Sanan-Mishra, 2021). When plants detect drought, transcription factors recognize and bind to specific DNA sequences, regulating the expression of downstream genes (Janiak *et al*., 2016; Baek *et al*., 2023). Accordingly, drought stress exerts a comprehensive impact on plants, extending beyond water deficit and encompassing intricate responses at the physiological, biochemical, and molecular levels.

The Assay for Transposase Accessible Chromatin with High-Throughput Sequencing (ATAC-seq), also known as the chromatin accessibility assay, is a critical tool in epigenomics (Yan *et al*., 2020). It enables the detection of accessible chromatin regions within cells, such as promoters, enhancers, and transcription factor binding sites, at high throughput (Li *et al*., 2019; Merrill *et al*., 2022). Well-established in animal research, ATAC-seq has successfully identified thousands of enhancers (Quillien *et al*., 2017; Bozek *et al*., 2019). However, its application in plants is limited, owing to the unique characteristics of plant cell walls, which resulted in fewer enhancers identified in the scant reports on plant research (Wang *et al*., 2022*a*; Lu *et al*., 2017). Multi-omics integrated analysis, on the other hand, is a promising new direction that combines individual omics results to explain their functions and mechanisms better comprehensively and systematically, addressing specific biological questions (Liu *et al*., 2023; Wang *et al*., 2022*b*; Chen *et al*., 2023). Luo *et al*. (Luo *et al*., 2022) discovered that combining ATAC-Seq with other genomic data, such as ChIP-Seq and RNA-Seq, can identify relevant cis-regulatory elements (CREs) in the biological genome. Consequently, this method is more potent for investigating gene functions, regulation, and expression mechanisms.

*Zoysia japonica* Steud., commonly referred to as zoysiagrass, is a perennial warm-season turfgrass from *Poaceae* family known for its strong resistance and wide-ranging applications (Guo *et al*., 2023; Wang *et al*., 2022*d*). It is used not only as a turfgrass for ornamental lawns and sports fields but also for soil conservation and landscaping, and it has a high economic value (Dong *et al*., 2021; Teng *et al*., 2021). Unfortunately, northern China experiences limited rainfall, requiring irrigation for the cultivation and maintenance of zoysiagrass. This leads to high water consumption in municipal landscapes. Previous research has confirmed inherent genotypic differences in the water demand mechanisms among different varieties of zoysiagrass. The discovery of drought-resistant genes in zoysiagrass can contribute to its well-established tissue culture and genetic transformation system. Zoysiagrass can serve as a reference species for improving turfgrass drought resistance in other species. Additionally, significant efforts by numerous scientists have yielded comprehensive data on the morphological and cytogenetic aspects of zoysiagrass (Tanaka *et al*., 2016; Wang *et al*., 2023). Multiple genes control drought resistance in zoysiagrass, and the physiological and biochemical processes it undergoes are regulated by various factors, including genetic, genetic interactions, and environmental factors (Xu *et al*., 2017; Feng *et al*., 2019). However, little is known about the genomics of drought resistance in zoysiagrass, which in turn limits progress in breeding efforts (Guan *et al*., 2022*b*; Cui *et al*., 2021). Therefore, it is critical to investigate drought resistance in zoysiagrass by exploring and utilizing resistance genes via multiomics technologies, thereby accelerating the genetic gain of key traits in zoysiagrass.

This study evaluated the drought resistance of zoysiagrass by determining physiological indicators under drought stress conditions to identify the optimal sampling time. Multiple omics techniques, including whole-genome sequencing (WGS), RNA sequencing (RNA-Seq), and assay for transposase-accessible chromatin sequencing (ATAC-Seq), were employed to analyze the mechanisms of drought stress. Candidate drought resistance genes were identified through transcriptome analysis, and their regulation by differential peaks in ATAC-Seq was investigated. The identified candidate genes were further validated using reverse transcription quantitative PCR (RT-qPCR). Additionally, motif analysis of the candidate genes led to the identification of upstream key regulatory transcription factors. This study focuses on discovering genes related to drought stress in zoysiagrass through multi-omics analysis, which provides a foundation for subsequent research on zoysiagrass genomics and molecular marker breeding.

## Materials and Methods

### Plant materials and growth conditions

The Turfgrass Research Institute at Beijing Forestry University provided the experimental materials, which were stored for 20 days at a 4 °C environment, including six zoysiagrass ecotypes: ‘Qingdao’ (QD), ‘Compadre’ (COM), ‘Zenith’ (ZEN), and ‘Meyer’ (M) as commercial ecotypes, and ‘X1’ and ‘X4’ as wild specimens collected from Northeast China.

The Turfgrass Research Institute at Beijing Forestry University provided stem segments with zoysiagrass sprouts, which were thoroughly washed and soaked in a nutrient solution for 3 hours. They were then immersed in a 0.15% FeSO_4_ solution for 1 hour before being rinsed three times with distilled water (Cai *et al*., 2015). Sterilized nutrient soil, sand, and vermiculite were mixed in a 2:1:1 ratio with sterilizing agents and placed in flowerpots before being moistened with water and set aside. Healthy stolon segments, each about 2 cm long at the base of the stem, were chosen, including one node with side shoots not yet emerging from the leaf sheath and side shoots about 1 cm on each side. These were planted in flowerpots, with eight stem segments of uniform size per pot. Six pots were prepared for each ecotype and placed in a growth chamber with a light cycle of 16 hours of light at 28 °C followed by 8 hours of darkness at 23°C, and the nutrient solution was kept at a weight of 400.0 g, indicating adequate moisture conditions.

### Experimental design for drought tolerance evaluation

Seedlings of zoysiagrass were grown for 70 days in a growth chamber with two biological replicates. Six ecotypes of zoysiagrass, each with two pots, were chosen for further testing in each experimental group. Control Group (CK), the photoperiod was programmed to be 16 hours of light at 28 °C and 8 hours of darkness at 23 °C. The soil weight was kept constant at 260.0 g, and the pot weight was 22.0 g. A nutrient solution was added, and the total weight was kept at 400.0 grams. Drought Stress Group (D), keeping all other conditions constant, the soil moisture content was kept at 10% by weight or 300.0 grams. Leaf tissue samples were collected for physiological parameter measurements on the 5th, 10th, 15th, 20th, and 25th days of treatment. Additionally, on the 15th day, ten mature and unfolded leaves were randomly selected from each pot for measuring leaf length and width. They were chosen as materials for RNA-seq and ATAC-seq based on the plants’ comprehensive drought tolerance assessment index.

The fresh weight (FW) of approximately 0.1 grams of fresh zoysiagrass leaves was weighed and recorded. A 10 mL centrifuge tube was filled halfway with water with the weighed needlegrass leaves, sealed, and set aside for 24 hours. After that, the needlegrass leaves were quickly removed, and the surface was blotted to remove any excess water. The saturated fresh weight (SFW) was noted thereafter. The recorded needlegrass leaves were placed in an envelope and blanched for 15 minutes at 105 °C. Afterward, the blanched needlegrass leaves were placed in a drying oven set to 65 °C and dried until they reached a constant weight (at least three days). The leaves were weighed again after drying and the weight was recorded as the dry weight (DW). The samples were kept to determine the soluble sugar content. The relative water content of the zoysiagrass leaves was calculated using the following formula as (FW-DW)/(SFW-DW) × 100%(Ma *et al*., 2018). Leaf proline content was determined as previously described (Bates *et al*., 1973).

### RNA extraction and ATAC libraries preparation

The RNeasy Plant Mini Kit from the QIAGEN company was used to extract RNA from zoysiagrass leaves. The experiment was carried out following the kit’s instructions. Novogene Technology Co., Ltd. performed sequencing after determining the concentration and quality of the RNA solution. We used FastQC software (version 0.11.9) to perform quality control on the raw sequencing data to assess and filter the sequencing data quality(Brown *et al*., 2017). We performed filtering and quality selection on the zoysiagrass transcriptome data. To ensure data quality, we used Trimmomatic with the following parameters to remove low-quality fragments:--phred33--LEADING:3--TRAILING:3--MINLEN:51 (Bolger *et al*., 2014). A second round of quality control was performed using FastQC to ensure the data’s accuracy. After quality control, the RNA-Seq data was aligned using HISAT2 software, and the results were saved in SAM format files(Kim *et al*., 2019). Picard’s SAMtools tool was used to compress and sort SAM-format files(Li *et al*., 2009). The completed data was saved in BAM format. The multiBam Summary command from deeptools (version 3.5.1) software was used to calculate the coverage of sequencing data in each interval (default 10 kb)(Ramirez *et al*., 2014). The plotCorrelation or plotPCA command was used to calculate the correlation of the transcriptome data in zoysiagrass.

The nuclear extraction reagent formulas of various *Gramineae* plants were used to extract nuclei and build libraries for ATAC-Seq from zoysiagrass (Qi *et al*., 2023). The Novoprotein Chromatin Profile Kit for Illumina extracted nuclei and fragment chromatin in zoysiagrass. The chromatin fragments were then purified with the Novoprotein Tagment DNA Extract Beads kit. The Novoprotein NovoNGS® Index Kit for Illumina® was used for PCR enrichment. The PCR products were purified, and the library size was determined using the Novoprotein DNA Clean Beads kit. Qubit was used to determine the concentration of the zoysiagrass library. An Agilent 2100 instrument was used for the initial library quality checks. Beijing Novogene Co., Ltd. then undertook the sequencing of high-quality zoysiagrass libraries.

### RT-qPCR validation in zoysiagrass

RNA was extracted from the leaves of the plant using the YEASEN MolPure® Plant Plus RNA Kit. The extracted RNA was then reverse transcribed into cDNA using the YEASEN Hifair® III 1st Strand cDNA Synthesis SuperMix for qPCR kit. The resulting cDNA templates from various zoysiagrass treatments were used for qPCR amplification experiments with qPCR primers (Supplementary Tab.2). As an internal reference, the Actin gene was used, with ACT-F primer sequence GGTCCTCTTCCAGCCATCCTTC and ACT-R primer sequence GTGCAAGGGCAGTGATCTCCTT. During the quantitative PCR amplification experiment, the Hieff® qPCR SYBR Green Master Mix kit from YEASEN was used to detect the expression levels of all candidate genes.

### RNA-seq and ATAC-seq data analysis

After obtaining the BAM files, the gene quantification analysis was done using the HTseq-count software (version 0.11.2)(Anders *et al*., 2015). Counting the number of reads within regions aligned to the reference genome based on gene positional information in the reference genome was required. DESeq2 (version 1.26.0) was used to statistically analyze the read count files to analyze gene expression further and perform functional enrichment(Love *et al*., 2014). Based on MapMan and Gene Ontology (GO) gene functional annotation information, a functional enrichment analysis of differentially expressed genes in zoysiagrass was performed. The zoysiagrass genome annotation contains gene functional annotation files for MapMan and GO. The statistical analysis of differential gene functional enrichment was performed in R using a two-tailed Fisher’s exact test based on the obtained data results to assess the enrichment level. *P* < 0.01 was the significance threshold for differentially expressed genes in zoysiagrass. R packages such as ggplot2, highcharter, and ggsci were used to create graphical representations of data. KEGG pathway enrichment analysis was performed on the identified genes using TBtools software(Chen *et al*., 2020), and the enriched pathways were visualized in R using the GGplot2 package. The zoysiagrass genome annotation contains KEGG pathway information.

Furthermore, the Institute of Lawns at Beijing Forestry University provided the ‘COM’ genome (Not publicly available yet, PRJNA1036829) for data alignment. All raw sequencing reads have been deposited in the NCBI Sequence Read Archive under project PRJNA1044241. ATAC-Seq data were aligned using BWA, based on alignment algorithms and Bowtie2(Li and Durbin, 2009). SAM (Sequence Alignment/Map) files were used to store the alignment results. The SAM format files were then compressed and sorted using the Picard toolkit’s SAMtools software(Li *et al*., 2009). The final organized data was saved in BAM (Binary Alignment/Map) format files for efficient storage and subsequent analysis. After filtering the zoysiagrass ATAC-Seq data, removing PCR duplicates with the “MarkDuplicates” feature in Picard version 2.20.3 software(Anxionnat *et al*., 2021), and excluding mitochondrial and chloroplast data with SAM tools(Li *et al*., 2009). MACS v2.1.2 software was used to identify chromatin-accessible regions in zoysiagrass and peak statistics(Zhang *et al*., 2008).

Differential peaks were identified using *P* < 0.01 as the significance threshold. Gene association and annotation were performed for significantly different peaks, focusing on the regions within 2 kb upstream and downstream of the transcription start site. The zoysiagrass genome FASTA file, annotated gene structure file, and BAM files generated by the analysis were imported into the Integrative Genomics Viewer (IGV) software for data visualization and motif prediction (Thorvaldsdottir *et al*., 2013). The MEME online tool was used to predict motifs, found at https://meme-suite.org/meme/tools/meme, and FigTree was used to visualize and analyze the evolutionary tree of target proteins(Bailey *et al*., 2009).

Two algorithms, Dynamic Tree Cut and Merged Dynamic, performed gene co-expression network analysis on 24 zoysiagrass samples to identify clustered gene groups with similar expression patterns(Langfelder *et al*., 2008). Previous studies have demonstrated that genes such as *OsDREB2A* (Dubouzet *et al*., 2003), *TaSnRK2.3* (Tian *et al*., 2013), *ZmVPP1* (Wang *et al*., 2016), *TaNAC2* (Mao *et al*., 2012), *OsNAC5* (Takasaki *et al*., 2010), *TaABF1* (Mao *et al*., 2022), and *HvWRKY38* (Xiong *et al*., 2010), among others, significantly enhance a plant’s drought tolerance. A homologous alignment was performed against the protein sequences of previously published drought-resistant genes (E-value less than 1e^-5^) to investigate further candidate genes related to drought stress, and homologous gene phylogenetic trees were constructed by using the IQ-TREE software(Lam-Tung *et al*., 2015).

## Results

### Drought tolerance assessment in zoysiagrasses

The relative water content of zoysiagrass leaves decreases continuously as the duration of drought stress increases (Fig. 1a). The relative water content of leaves in X1 zoysiagrass begins to fall sharply on the 5th day and continues until the 20th day when there is little change. In the case of X4 zoysiagrass, the relative water content of the leaves begins to decline on the fifth day and continues to fall sharply until the 25th day, when it reaches a significantly lower level.

**Fig.1.**
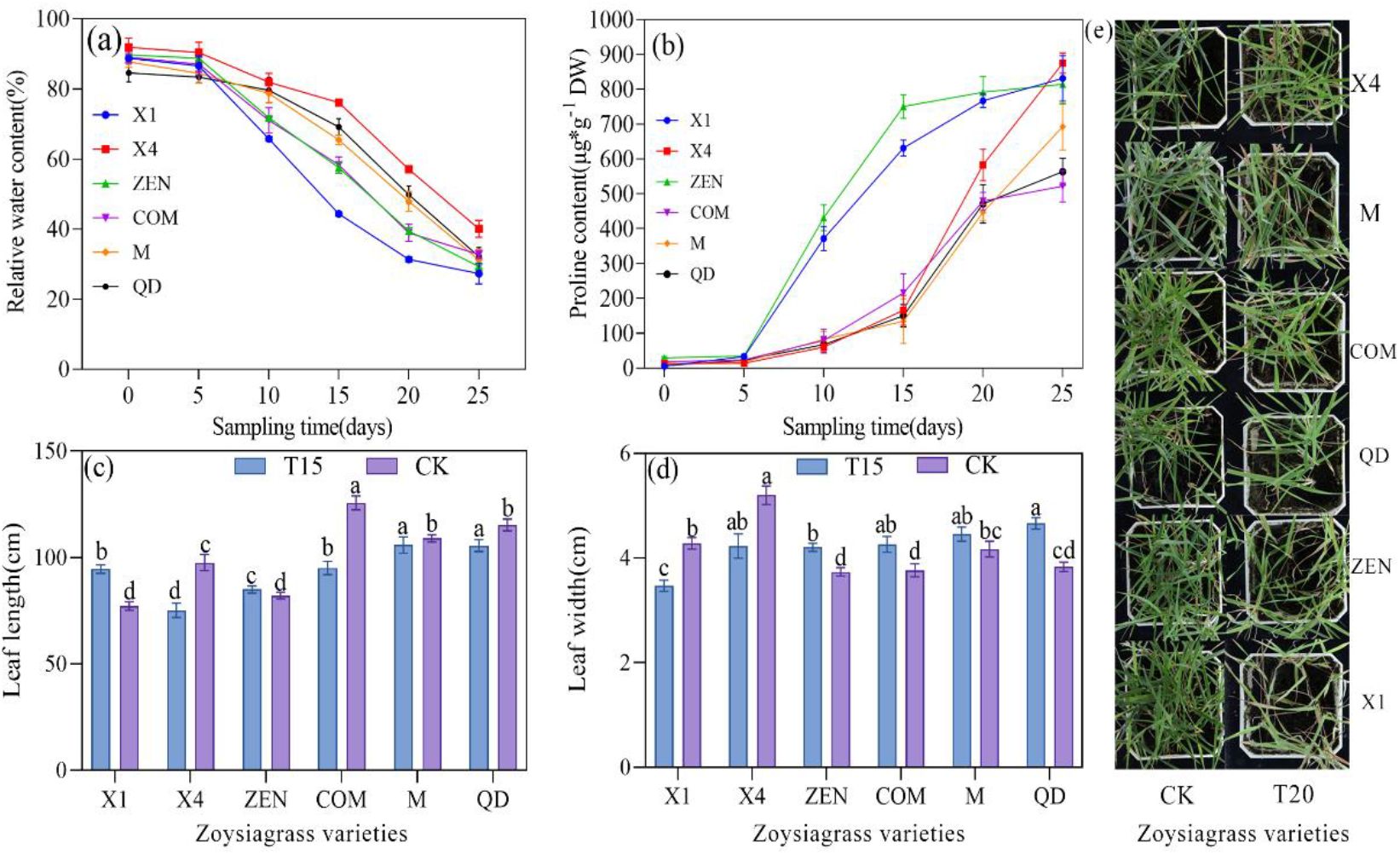
Physiological indicators and phenotypic growth in zoysiagrass. (a)Relative water content in zoysiagrass leaf. (b)Proline content in zoysiagrass leaf. (c) and (d) Comparison of phenotypic traits under drought stress after 15 days of growth. (e) Comparison of drought stress phenotype traits after 20 days of growth. T15, 15 days of zoysiagras growth under drought stress treatment. T20, 20 days of zoysiagrass growth under drought stress treatment. CK, Control group. The same letters in the same color bar chart showed no significant difference (*P* > 0.05), the adjacent letters showed significant difference (*P* < 0.05), and the separated letters showed significant difference. Error bars indicate the standard error of the mean.

The proline content of zoysiagrass leaves steadily increases as the duration of drought stress increases (Fig. 1b). In the case of ZEN zoysiagrass, the proline content in the leaves begins to rise sharply on the 5th day and continues to rise until the 20th day, when there is little change. Starting on the 15th day, the proline content in leaves of M, QD, and X4 zoysiagrass increases dramatically. Based on the proline content trend and proline content on the 20th day, the preliminary drought tolerance ranking of zoysiagrass was established: X4 > M > COM > QD > ZEN > X1.

Significant differences in zoysiagrass phenotypes emerged after the 15th day of drought treatment. X1, M, and ZEN zoysiagrass leaves began to wilt by the 20th day, with X1 zoysiagrass exhibiting the most significant leaf wilting, and dry grass appeared beneath the plants (Fig. 1c). The length of zoysiagrass leaves was slightly shorter than that of the control group on the 15th day of drought treatment, and the width was slightly wider than that of the control group (Fig. 1d).

The subordination function method assessed the drought tolerance of six zoysiagrass ecotypes (Tab. 1). Plant resistance is positively correlated with subordination functions. Based on the average subordination function values, the drought tolerance ranking of zoysiagrass is as follows: X4 > M > COM > QD > ZEN > X1.

**Tab.1.** Evaluation of drought resistance of zoysiagrass.

| Ecotype | Relative Water Content | Proline Subordination Value | Average Subordination Value | Resistance Ranking |
| --- | --- | --- | --- | --- |
| X1 | 0 | 0.194 | 0.097 | 6 |
| X4 | 1.000 | 0.949 | 0.974 | 1 |
| ZEN | 0.317 | 0 | 0.158 | 5 |
| QD | 0.296 | 0.868 | 0.582 | 4 |
| M | 0.646 | 1.000 | 0.823 | 2 |
| COM | 0.369 | 0.878 | 0.623 | 3 |

### Assessment of RNA-seq and ATAC-seq data

The extracted nuclei from 24 zoysiagrass samples (control and drought-treated groups) were assessed to obtain high-quality ATAC-seq sequencing data for zoysiagrass. The extracted nuclei were round and intact (Supplementary Fig.S1A) with minimal breakage (Supplementary Fig.S1B), indicating good nuclear quality suitable for library construction. The COM zoysiagrass in the control group served as an example of how to evaluate the constructed libraries after library construction. The library insert size and peak values were appropriate for subsequent sequencing, indicating high library quality (Supplementary Fig.S1C&D). The detection results confirmed the data’s dependability, making it appropriate for the subsequent discovery of resistance genes in zoysiagrass.

The raw RNA-Seq data from 24 zoysiagrass samples was filtered to obtain high-quality sequencing data. The transcriptome data were then aligned to the assembled COM zoysiagrass genome using the HISAT2 software. Following alignment, data reproducibility was evaluated, and anomalies and outliers in the data were identified and removed using two methods, Relative Log Expression (RLE) and Variance Stabilizing Transformation (VST), to ensure the data’s accuracy and reliability (Supplementary Fig.S2). The results consistently showed a strong correlation within the same treatment group and significant differences between treatments, indicating the data’s reliability, which can be used to discover resistance genes in zoysiagrass in the future.

### Identification of drought-tolerant genes

The gene co-expression and clustering analysis network revealed 24 modules or classes used to build an evolutionary tree and generate an expression heatmap (Fig.2). The MEbrown, MEtan, MEsalmon, and MElightyellow gene modules in zoysiagrass showed enriched regulation after drought treatment (Fig.2C). To elucidate the regulatory mechanisms further, gene ontology (GO) analysis of the 24 modules revealed enrichment in 108 pathways, and 35 pathways were linked to drought resistance(). A combined analysis of GO enrichment results and regulatory modules revealed that MEbrown, MEtan, MEsalmon, and MElighyellow modules were enriched in processes such as “cellular response to water deprivation,” “very long-chain fatty acid metabolic process,” “cuticle development,” and the “abscisic acid-activated signaling pathway.” These biological processes are highly relevant to drought stress, and their enrichment in gene modules confirms their involvement in zoysiagrass drought regulation.

**Fig.2.**
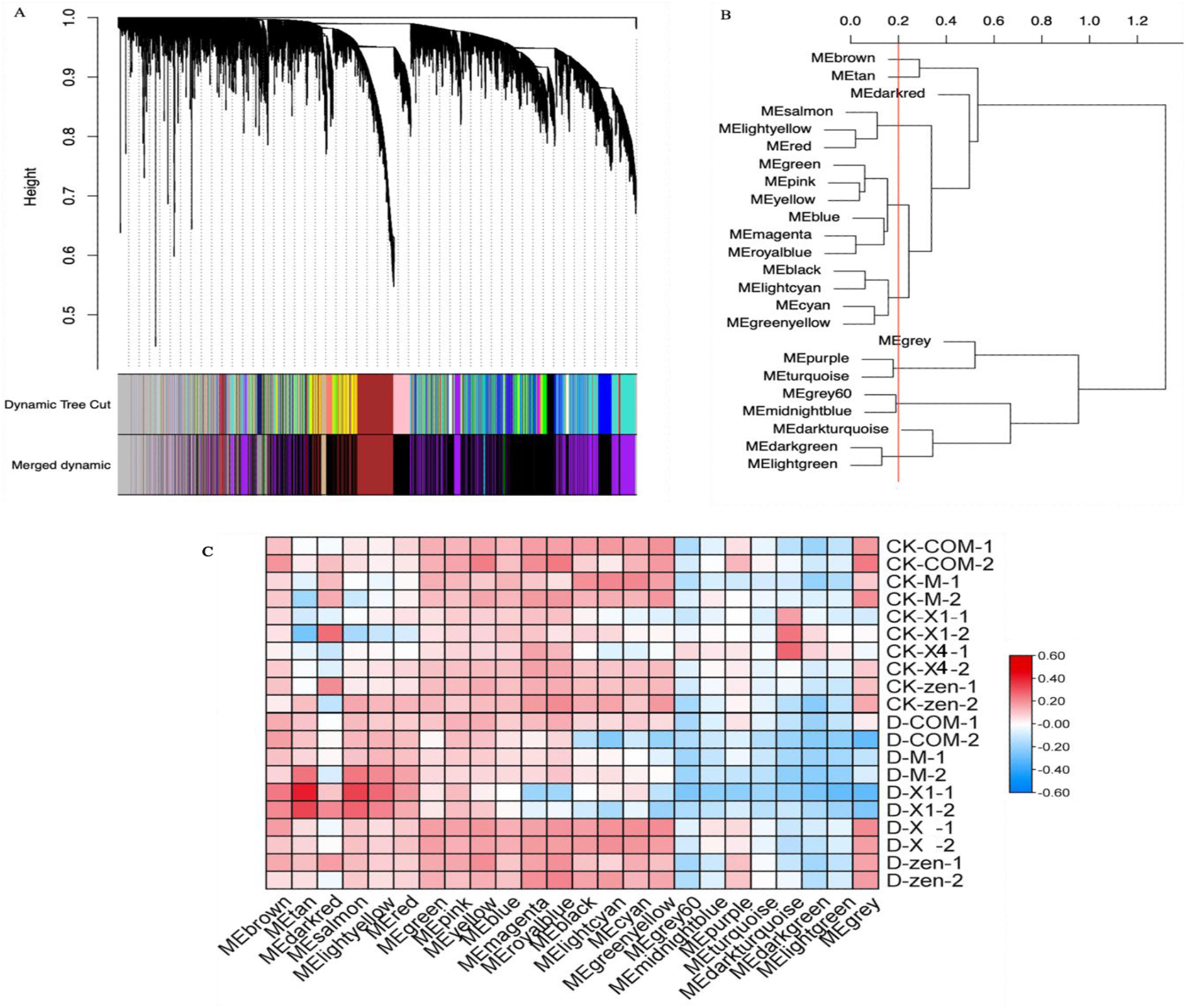
Expressed genes in zoysiagrass. (A)Zoysiagrass gene expression and clustering. (B) Module evolutionary tree analysis. (C) Gene co-expression heatmap of the modules.

ATAC-Seq sequencing and analysis were performed on 24 samples (12 samples with two replicates) to investigate the identified resistance genes further and obtain the conserved chromatin-accessible regions in zoysiagrass. Each sample was sequenced and analyzed at a depth of 10 GB, corresponding to approximately 19x coverage depth (Supplementary Tab.1). Following initial filtering of the zoysiagrass ATAC-Seq raw data, the Bowtie2 software aligned to the assembled zoysiagrass genome of the COM ecotype. To ensure reliability and validity, a repeatability test was performed on the zoysiagrass ATAC-Seq data. The findings revealed strong data correlation within the same treatment groups and significant differences between treatment groups. It demonstrates the data’s dependability for future research into resistance genes in zoysiagrass (Supplementary Fig.S4). Distribution statistics of peak regions in zoysiagrass were calculated to investigate the impact of treatments on gene functionality (Supplementary Fig.S5). Peak expression differences were used to identify differential peaks with significant differences in expression (Fig. 3A). Gene association and annotation related to differentially expressed genes’ peak regulation were performed. The positional information of differential peaks was used to identify 1,694 genes regulated by differential peaks and related to drought treatment (Fig. 3B).

**Fig.3.**
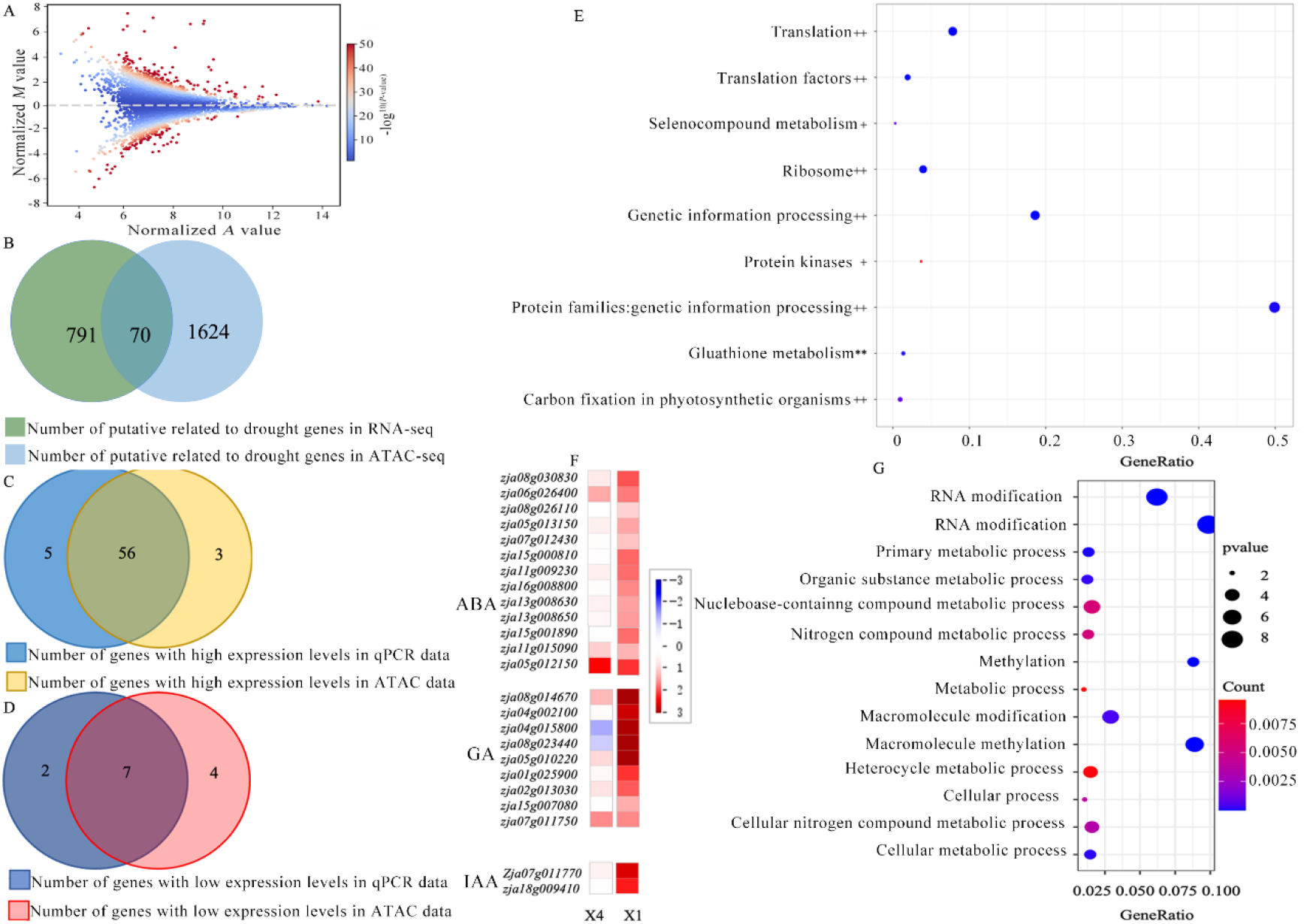
Differential expressional genes analysis of zoysiagrass. (A) Differential peak analysis of zoysiagrass. (B) Drought-related genes in zoysiagrass. (C) and (D) Drought-related genes in zoysiagrass. (E) * indicates pathways enriched for Class Ⅰ genes, ** indicates pathways enriched for Class Ⅱ genes, + indicates pathways enriched for Class Ⅲ genes, ++ indicates pathways enriched for Class Ⅳ genes. (F) KEGG analysis of zoysiagrass. (G) GO analysis of zoysiagrass.

A joint analysis was performed with high confidence to investigate drought-tolerant genes further, combining the drought-related genes identified in the transcriptome with the genes regulated by differential peaks found in the ATAC-Seq data. This study yielded a list of 70 genes linked to drought stress (Fig.3B, Supplementary Tab.2).

The discovered genes were subjected to RT-qPCR validation to further validate the reliability of the genes identified in this study. The 2^-ΔΔCt^ method was used to calculate the relative gene expression levels. The RT-qPCR results were generally consistent with the trends observed in the transcriptome data (Supplementary Tab.2, Fig. 3C, D). Sixty-three high-confidence genes associated with drought stress were identified (Supplementary Fig.S6).

To investigate the expression patterns of these genes in zoysiagrass, their expression levels were analyzed, and genes were classified into four classes based on gene expression level correlation. Class I and II genes were downregulated in zoysiagrass, while Class III and IV genes were upregulated to varying degrees (Fig. 4).

**Fig.4.**
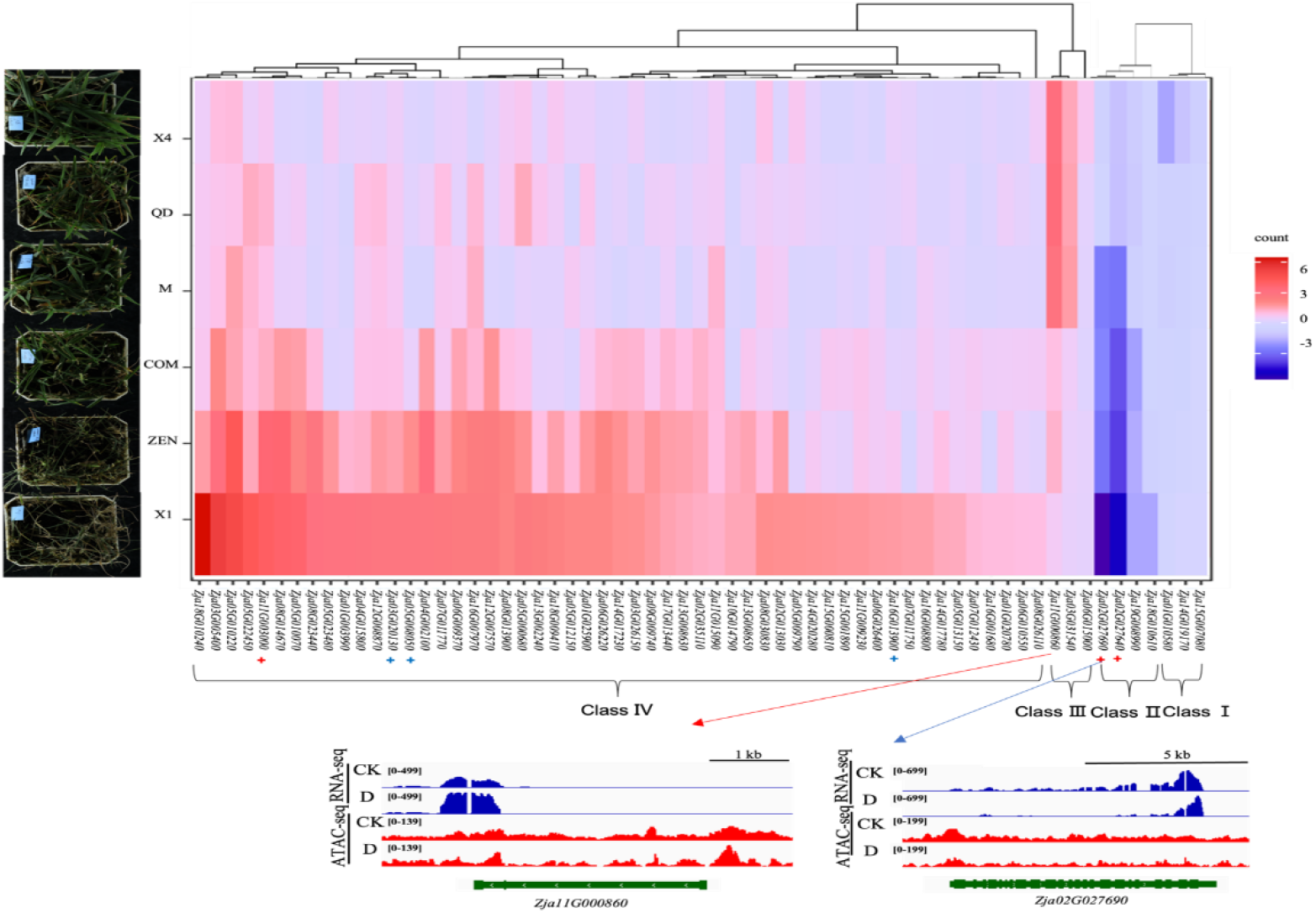
Expression of drought-resistant candidate genes in zoysiagrass. CK is the experimental control group, D is the drought treatment group, red + markers are expansion gene markers, and blue + markers are positive selection gene markers.

According to a transcriptomics and epigenomics integration analysis, the intensity of ATAC-Seq peaks is positively correlated with gene expression levels. The zoysiagrass gene *Zja11G000860*, for example, was highly expressed in drought-stressed zoysiagrass, and its ATAC-Seq peak intensity was higher than that of the CK zoysiagrass gene expression. In contrast, drought stress inhibited the expression of the zoysiagrass gene *Zja02G027690*, and its ATAC-Seq peak intensity was lower than that of the CK zoysiagrass gene expression (Fig. 4).

Furthermore, the expansion genes *Zja02G027640* and *Zja02G027690* were found to have downregulated expression in zoysiagrass of various ecotypes, and their expression levels decreased as zoysiagrass drought tolerance decreased. The expansion gene *Zja11G003000*, as well as the positively selected genes *Zja05G008050* and *Zja03G020130*, showed upregulated expression in zoysiagrass of various ecotypes, with expression levels increasing as zoysiagrass drought tolerance decreased (Fig. 4).

According to phylogenetic trees of homologous genes, the *Zja03G031540* gene in zoysiagrass shares high homology with the *SnRK2.3* gene. Furthermore, the zoysiagrass gene *Zja11G000860* shares high homology with the *NAC2* and *NAC5* genes (Supplementary Fig.S7). The zoysiagrass genes *Zja03G031540* and *Zja11G000860* have high homology with previously identified drought-resistant genes. They play a role in drought tolerance in zoysiagrass and are essential genes for improving drought resistance.

The differential expression of these genes in zoysiagrass was examined to confirm whether they are associated with drought stress in zoysiagrass. The genes *Zja03G031540* and *Zja11G000860* were found to be significantly upregulated in X4, M, and QD zoysiagrass, while they were found to be significantly downregulated in COM, ZEN, and X1 zoysiagrass. This expression pattern corresponds to the phenotypic findings, indicating that *Zja03G031540* and *Zja11G000860* are candidate genes related to drought stress in zoysiagrass (Fig.4, Supplementary Fig.S8).

### Analysis of drought-tolerant candidate gene’s function

The candidate genes were enriched in genetic information processing, protein families related to genetic information processing, translation, ribosome, and mitochondrial biogenesis pathways using KEGG (Fig. 3E&F). These pathways are critical for protein synthesis, which plants need to regulate and adapt to environmental changes when subjected to drought stress.

Expansion genes *Zja02G027690* and *Zja02G027640* related to drought stress were found to be enriched in the glutathione metabolism pathway, contributing to zoysiagrass drought resistance. When plants are subjected to drought or salt stress, glutathione regulates the cellular redox balance. Positively selected genes with high relevance to zoysiagrass evolution include *Zja16G013900*, *Zja03G020130*, and *Zja05G008050*, which are enriched in genetic information processing and protein families related to genetic information processing pathways. Additionally, *Zja11G003000*, Zja03G020130, and Zja05G00805 genes were enriched in metabolic functions, confirming their high relevance to stress tolerance (Fig.4, Fig.3 E, F&G). Furthermore, expansion genes *Zja02G027690* and *Zja02G027640* in class II were enriched in the glutathione metabolism pathway, while class III and IV genes were enriched in stress-related pathways. *Zja03G031540* and *Zja11G000860*, class III genes, were enriched in Selenium compound metabolism and Protein kinase pathways, indicating that zoysiagrass responds to environmental stress by regulating stress responses and antioxidant defenses. Genes *Zja03G031540* and *Zja11G000860* were more expressed in drought-tolerant X4 zoysiagrass than in less drought-tolerant X1 zoysiagrass.

Class IV zoysiagrass genes, including *Zja08G030830*, *Zja06G026400*, *Zja08G026110*, *Zja05G013150*, *Zja07G012430*, *Zja15G000810*, *Zja11G009230*, *Zja16G008800*, *Zja13G008630*, *Zja13G008650*, *Zja15G001890*, *Zja11G015090*, and *Zja05G012150*, were enriched in the abscisic acid (ABA) pathway. Genes Zja08G014670, *Zja04G002100*, *Zja04G015800*, *Zja08G023440*, *Zja05G010220*, *Zja01G025900*, *Zja02G013030, Zja15G007080*, and *Zja07G011750* were enriched in the gibberellin (GA) pathway. Genes *Zja07G011770* and *Zja18G009410* were enriched in the indole acetic acid (IAA) pathway. The expression of these genes showed variation among different between zoysiagrass varieties, which is linked to their drought resistance (Fig.3E&F).

A GO enrichment analysis was performed to further elucidate these candidate genes’ functions, revealing that these genes are primarily enriched in metabolic processes and methylation functions in the biological process category (Fig. 3G). The mechanism of gene regulation in response to drought stress in zoysiagrass was further elucidated using motif analysis, and the motifs of 6 high-confidence genes were identified (Fig.5). These motifs were similar to previously published *ERF/DREB* (O’Malley *et al*., 2016) and *C2H2* (Laity *et al*., 2001).

**Fig.5.**
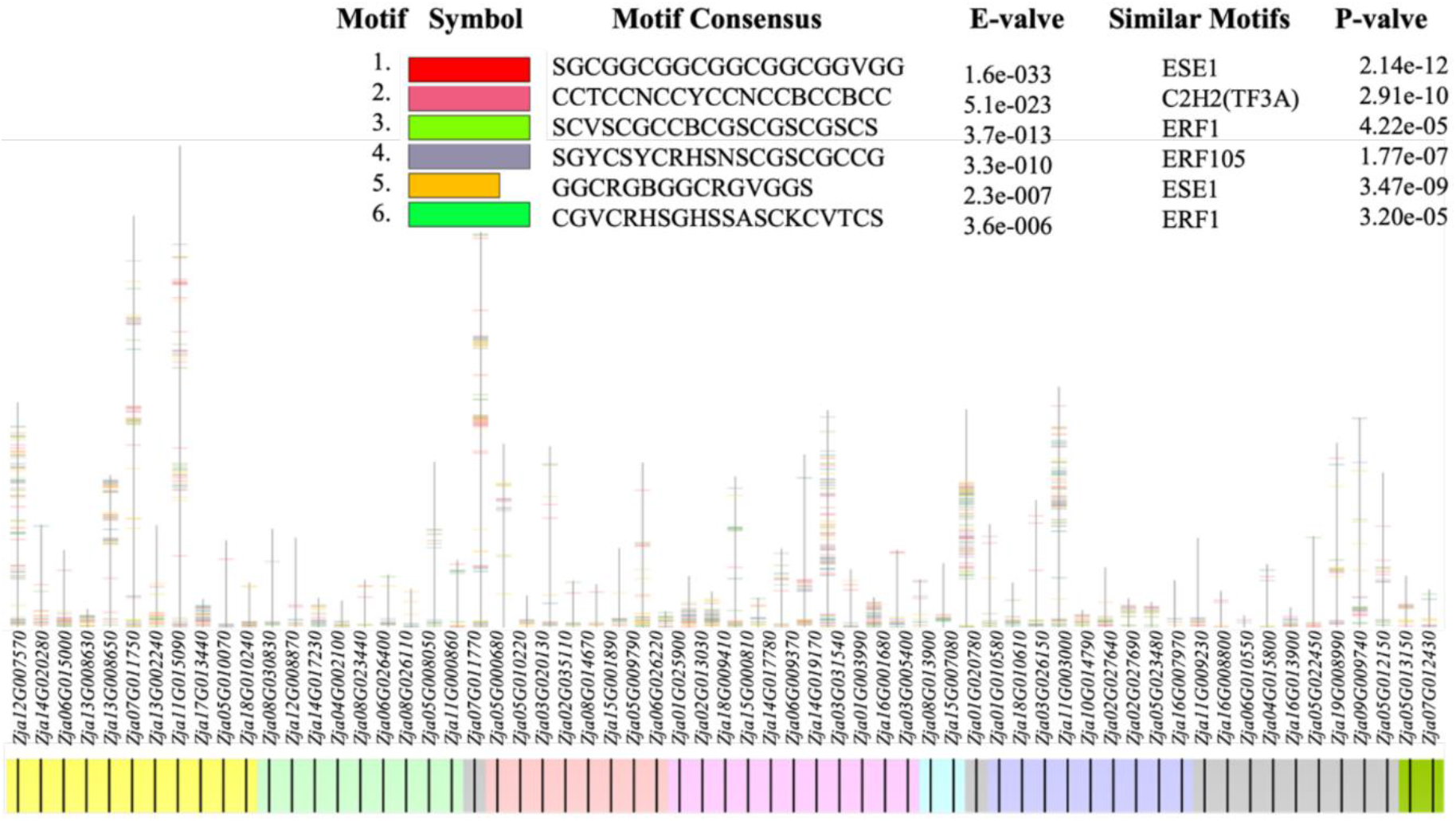
Motif analysis of drought-resistant candidate genes in zoysiagrass.

## Discussion

In regions of northern China, where warm-season lawn grasses are cultivated, irrigation is necessary, posing challenges to lawn maintenance standards. By improving the drought tolerance of these lawn grasses, it is possible to reduce irrigation requirements while preserving the visual appeal of lawns, which may lead to a more efficient utilization of valuable resources, including labor, materials, and finances. In our study, the discovery of drought-resistance genes in zoysiagrass holds implications beyond enhancing genetic gain for drought-resistance traits.

Drought tolerance in zoysiagrass was evaluated using experiments and physiological parameter measurements to determine the duration of drought treatment. On the 15th day of drought stress, the water and proline content of zoysiagrass rapidly changed, possibly in response to the drought stress. By the 20th day of treatment, the observed phenotypes of zoysiagrass aligned with the experimental data. X1 and ZEN exhibited low leaf water content and high proline content, resulting in varying degrees of leaf curling and slightly increased width compared to the control group. When exposed to drought, zoysiagrass exhibits relatively late phenotypic changes related to its inherent drought tolerance. The experiments discovered that after 35 days of drought stress, zoysiagrass could recover to some extent when rewatered, except for X1 and ZEN zoysiagrass, which could not fully recover. Finally, the drought resistance of different zoysiagrass ecotypes varies. These differences can be used to discover relevant genes in zoysiagrass, which may be various transcription factors are involved in the regulation of stress-inducible genes.

The candidate genes’ expression levels were examined to investigate their expression patterns. It was discovered that there is a link between gene expression levels and zoysiagrass phenotypes. Among the high-confidence genes related to drought stress discovered in this study, *Zja08G030830*, *Zja06G026400*, *Zja08G026110*, *Zja05G013150*, *Zja07G012430*, *Zja15G000810*, *Zja11G009230*, *Zja16G008800*, *Zja13G008630*, *Zja13G008650*, *Zja15G001890*, *Zja11G015090*, and *Zja05G012150* are enriched in the ABA pathway. The metabolism of ABA and the response of plants to drought stress are inextricably linked. It has been reported that the ABF1 gene regulates plant hormone ABA signaling and that ABF1 gene expression can improve drought resistance in wheat and Arabidopsis (Xu *et al*., 2008). Further validation and utilization of candidate genes enriched in the ABA pathway may be beneficial for improving drought resistance in zoysiagrass.

Drought stress can be mitigated by regulating growth hormone levels in zoysiagrass. On the one hand, Drought stress causes gene enrichment in the mitochondrial biogenesis pathway, which provides more energy to adapt to different growth conditions and stages. In response to drought stress, zoysiagrass has highly regulated gene expression and biological function, as evidenced by the enrichment of these gene pathways, which aid in zoysiagrass’s response to external drought stress. On the other hand, the enrichment of zoysiagrass genes in the functions of RNA methylation and RNA modification suggests that genes are regulated in response to environmental stress. Enrichment in primary, cellular, and organic substance metabolic processes also suggests active gene expression in metabolic pathways, which may means that zoysiagrass synthesizes energy and organic compounds in response to environmental changes to sustain life. Furthermore, enrichment in nitrogen compound metabolic processes, nucleobase-containing compound metabolic processes, heterocycle metabolic processes, cellular aromatic compound metabolic processes, organic cyclic compound metabolic processes, and RNA metabolic processes emphasizes the importance of these genes in protein and nucleic acid synthesis, which is critical for the response of zoysiagrass to external drought stress.

The *AP2/ERF* superfamily includes the Ethylene Response Factor (*ERF*)/Dehydration-Responsive Element-Binding (*DREB*) and Zinc Finger transcription factor domains (Mizoi *et al*., 2012; Lata *et al*., 2014). These transcription factors help plants respond to biotic and abiotic stressors like drought and low and high temperatures. *DREB* is a transcription factor family associated with dehydration. During dehydration stress, this transcription factor family regulates gene expression by binding to specific DNA sequences (such as ABA) to assist plants in adapting to environmental stress (Je *et al*., 2014; Dubouzet *et al*., 2003). In summary, the drought-related genes in zoysiagrass are high-confidence genes with motifs similar to *ERF/DREB* and Zinc Finger transcription factor domains, indicating their role in responding to environmental stress, particularly drought. The discovery of motifs that resemble the *ERF* (O’Malley *et al*., 2016) and *C2H2* (Laity *et al*., 2001) transcription factor family domains and influence the expression of candidate genes provides a new direction and resource for investigating zoysiagrass’ drought resistance mechanism. The discovery of key motifs and candidate genes lays the groundwork for improving zoysiagrass drought tolerance. Further validation and utilization of these genes are critical for improving drought resistance in zoysiagrass.

### Conclusion

Drought tolerance assessments for six ecotypes were conducted using physiological indicators of zoysiagrass. Sixty-three high-confidence genes related to drought stress and six motifs involved in drought regulation were identified using integrated omics analysis and RT-qPCR validation methods. The analysis revealed a positive relationship between ATAC-Seq peak intensity and gene expression levels in zoysiagrass. The expression of high-confidence genes is linked to resistance assessments and phenotypes in zoysiagrass. According to the findings, the newly discovered resistance genes play a role in actively regulating gene responses to external drought stress pressure.

## Supplementary data

Fig.S1. Extracting nuclei and checking library in zoysiagrass.

Fig.S2. Quality for RNA-seq data in zoysiagrass.

Fig.S3. Enrichment analysis of zoysiagrass.

Fig.S4. Quality for isolation and ATAC-seq data in zoysiagrass.

Fig.S5. Distribution of peak in gene functional elements.

Fig.S6. Drought-resistant candidate genes in zoysiagrass.

Fig.S7. Evolutionary tree of homologous genes.

Fig.S8. The drought response of zoyssiagrass.

Tab.1. Data and analysis of ATAC-seq in zoysiagrass.

Tab.2. Drought-related genes and its information of RT-qPCR in zoysiagrass.

## Author contributions

L. S. and Z. Q. conceived and drafted the manuscript. S.L., Z.Q., M.W., J. Z., and L. H. analyzed the data. J. C. and L. X. provided the plant resources. L. H. and Y. C. contributed to the conception of the study and finalized the manuscript. All authors read and approved the final manuscript.

## Conflict of interest

No conflict of interest declared.

## Funding

This study was supported by the National Key R&D Program, National Natural Science Foundation of China (31971770).

## Data availability

All raw data were deposited in the Genebank Short Read Archive (https://submit.ncbi.nlm.nih.gov/subs/sra/SUB13996434, https://www.ncbi.nlm.nih.gov/sra/PRJNA1044241)

## Notes

### Competing Interest Statement

The authors have declared no competing interest.

