## Supplementary data for "Integration of ATAC-seq and RNA-seq identifies novel candidate genes associated with drought tolerance in *Zoysia japonica* Steud"

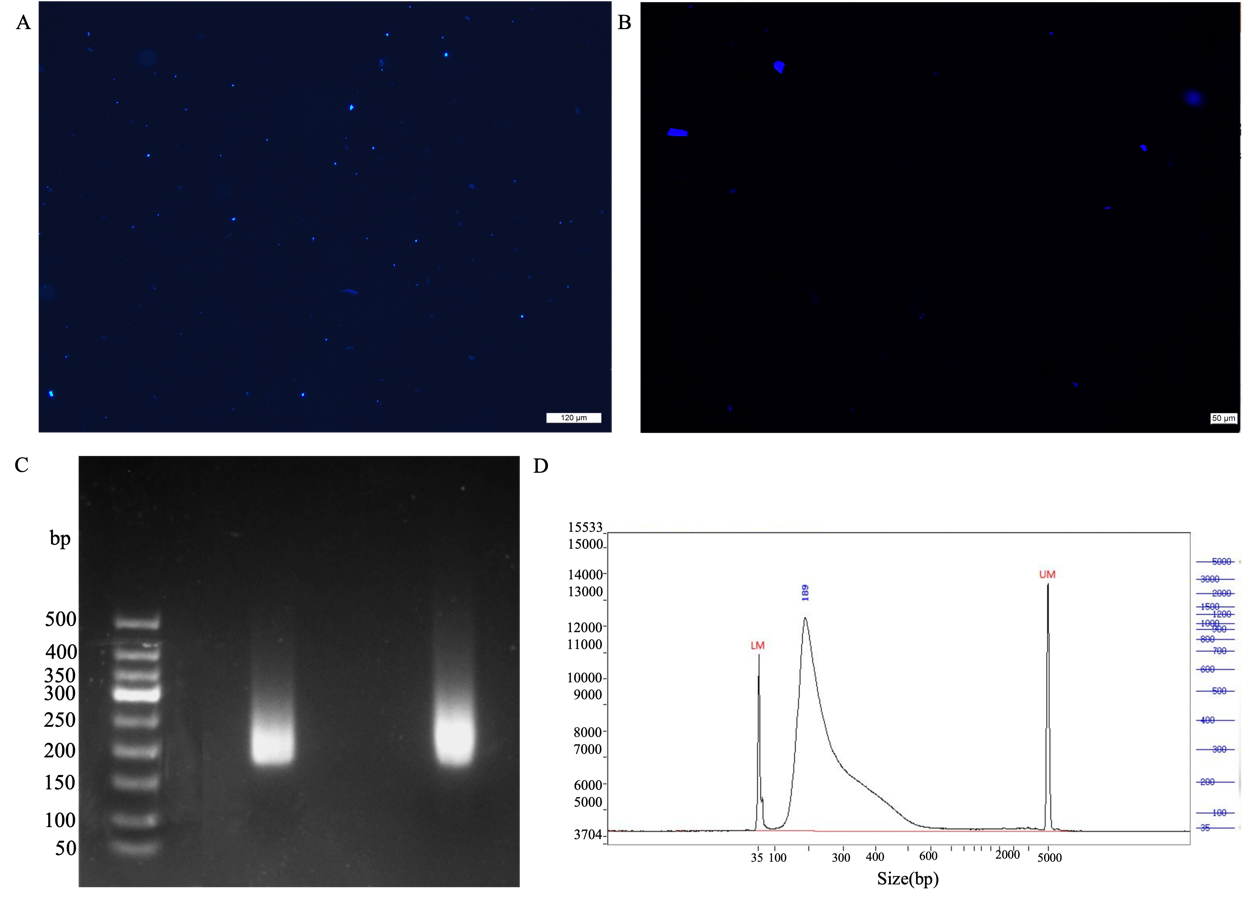

**Fig.S1.** Extracting nuclei and checking library in zoysiagrass. (A) Extraction of nuclei from zoysiagrass leaves. (B) Ethidium bromide-stained gel for checking nuclei. (C) Agarose gel electrophoresis of the library. (D) Library quality check.

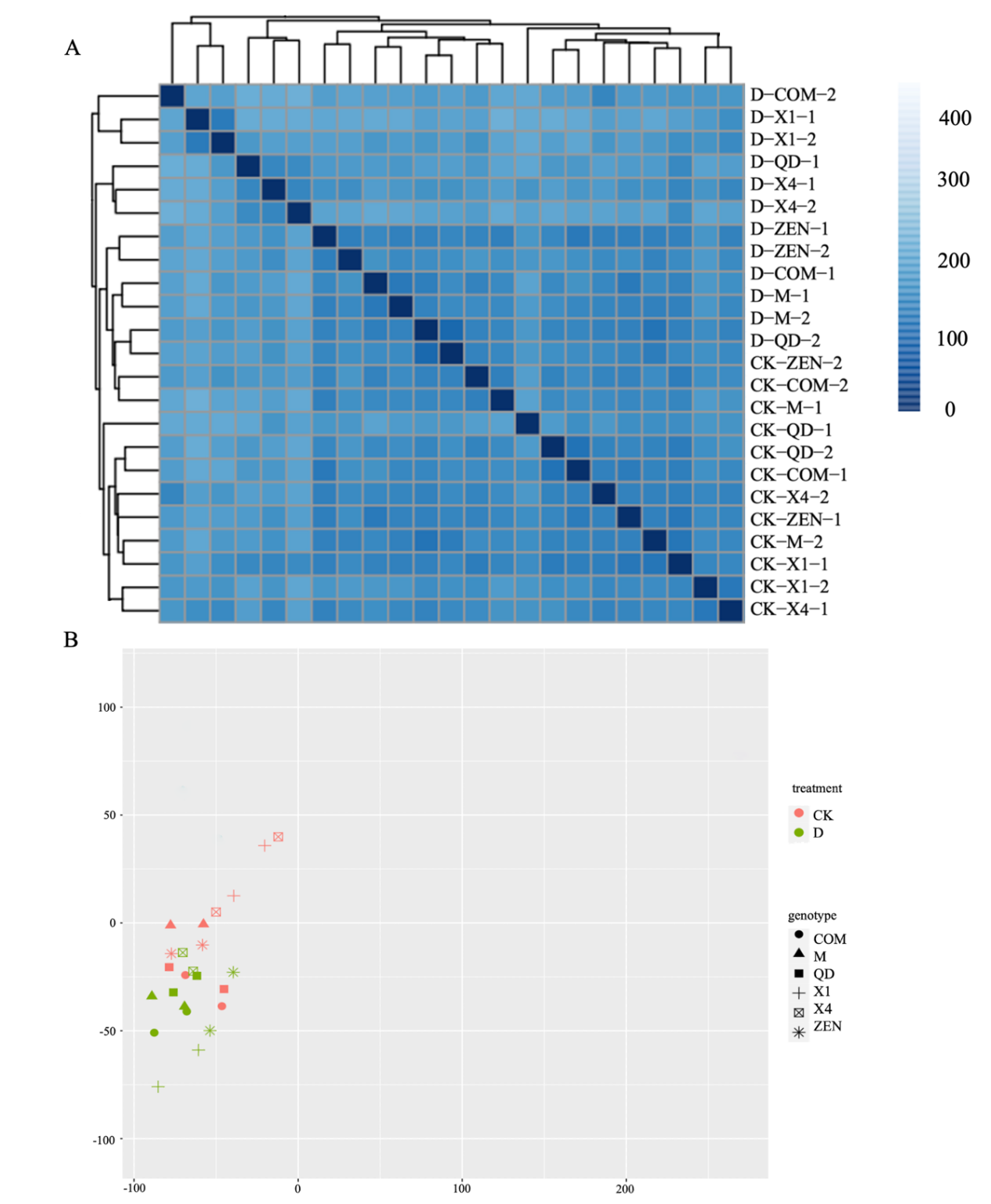

**Fig.S2.** Quality for RNA-seq data in zoysiagrass. (A) Reproducibility analysis of Zoysiagrass RNA-seq data using the Relative Log Expression (RLD) method. (B) Reproducibility analysis of Zoysiagrass RNA-seq data using the Variance Stabilizing Transformation (VSD) method.

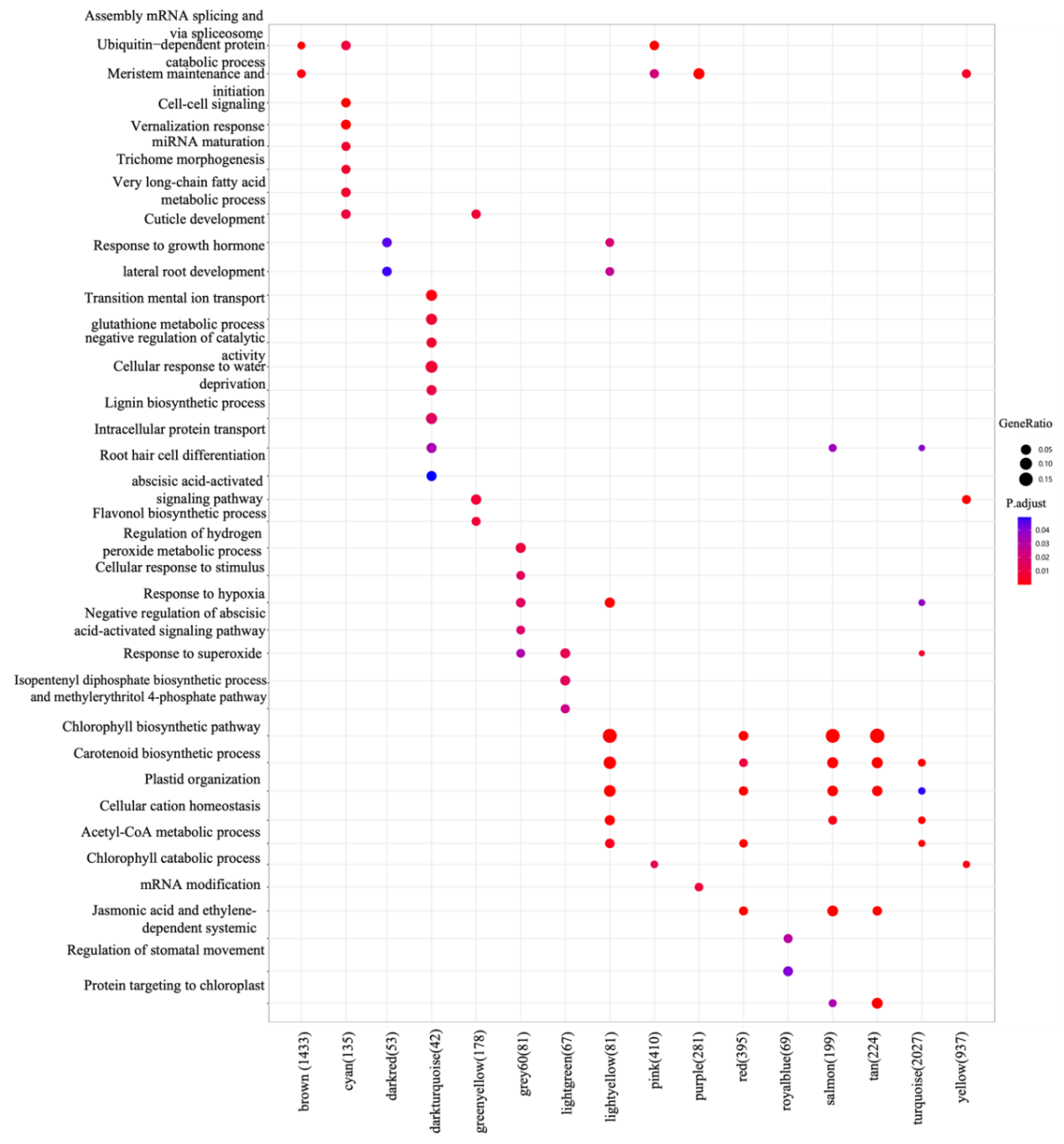

**Fig.S3.** Enrichment analysis of zoysiagrass.

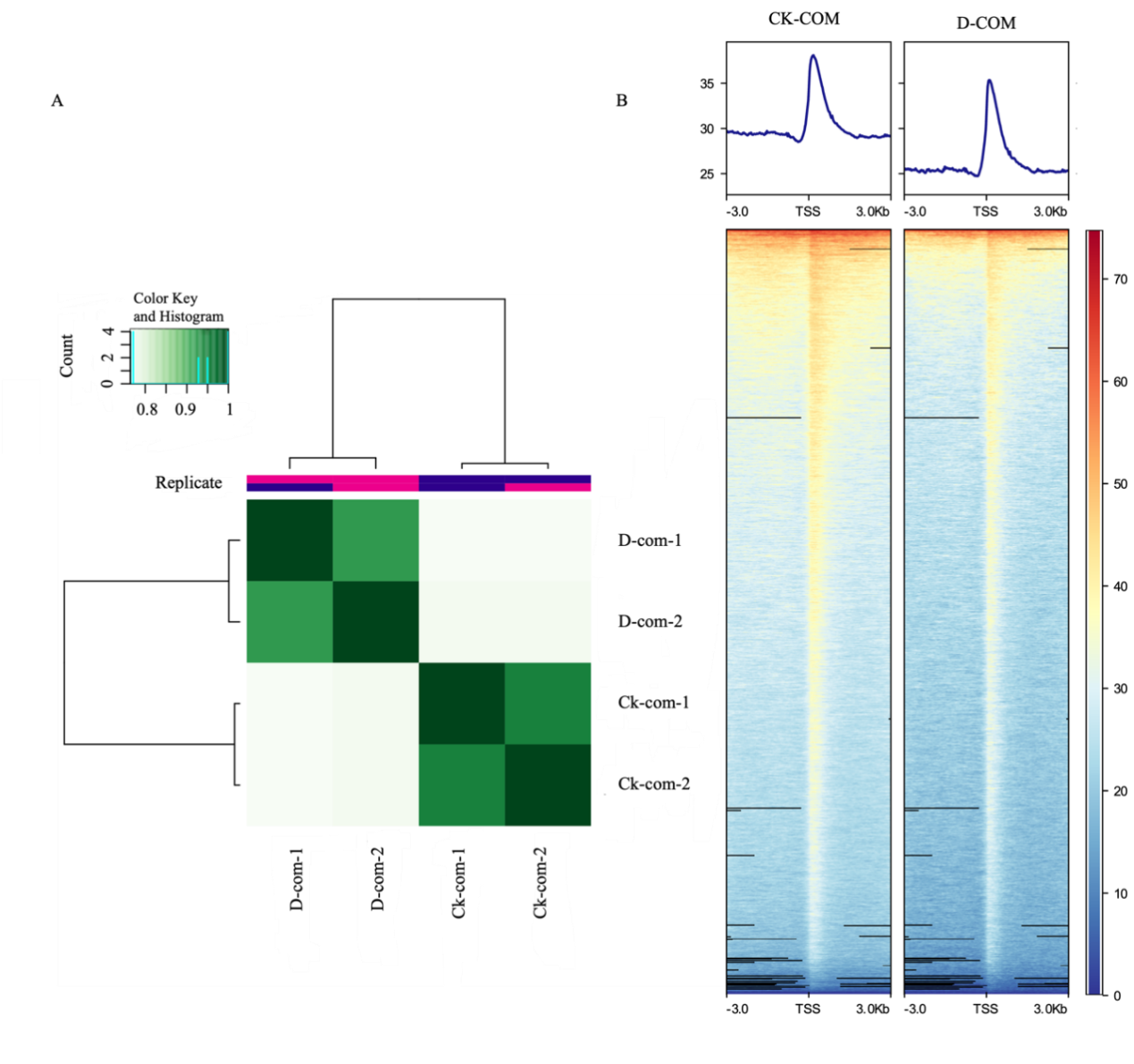

**Fig.S4.** Quality for isolation and ATAC-seq data in zoysiagrass. (A): Reproducibility assessment of Zoysiagrass ATAC-seq data. (B): Signal plots for calling peaks.

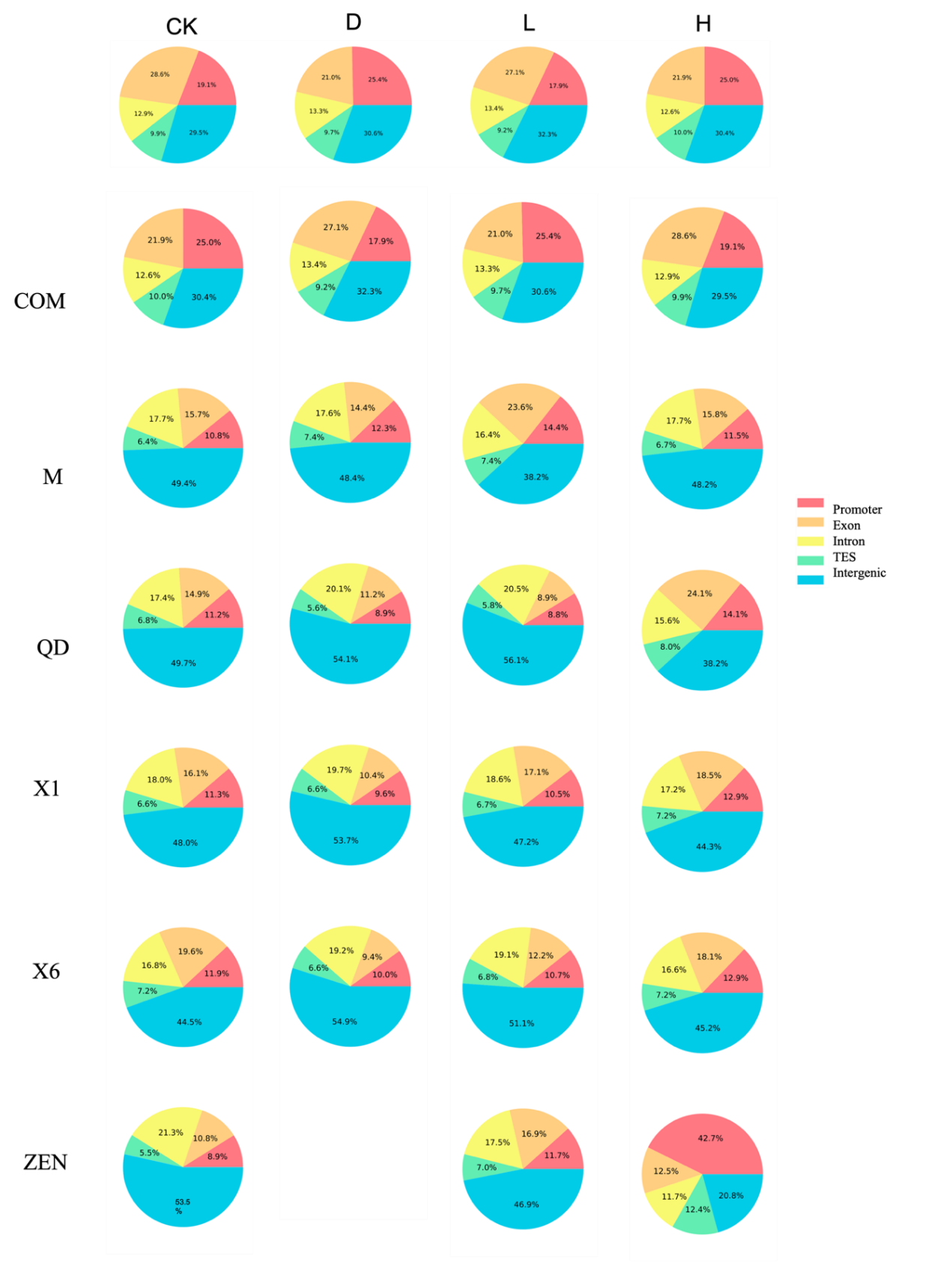

**Fig.S5.** Distribution of peak in gene functional elements.

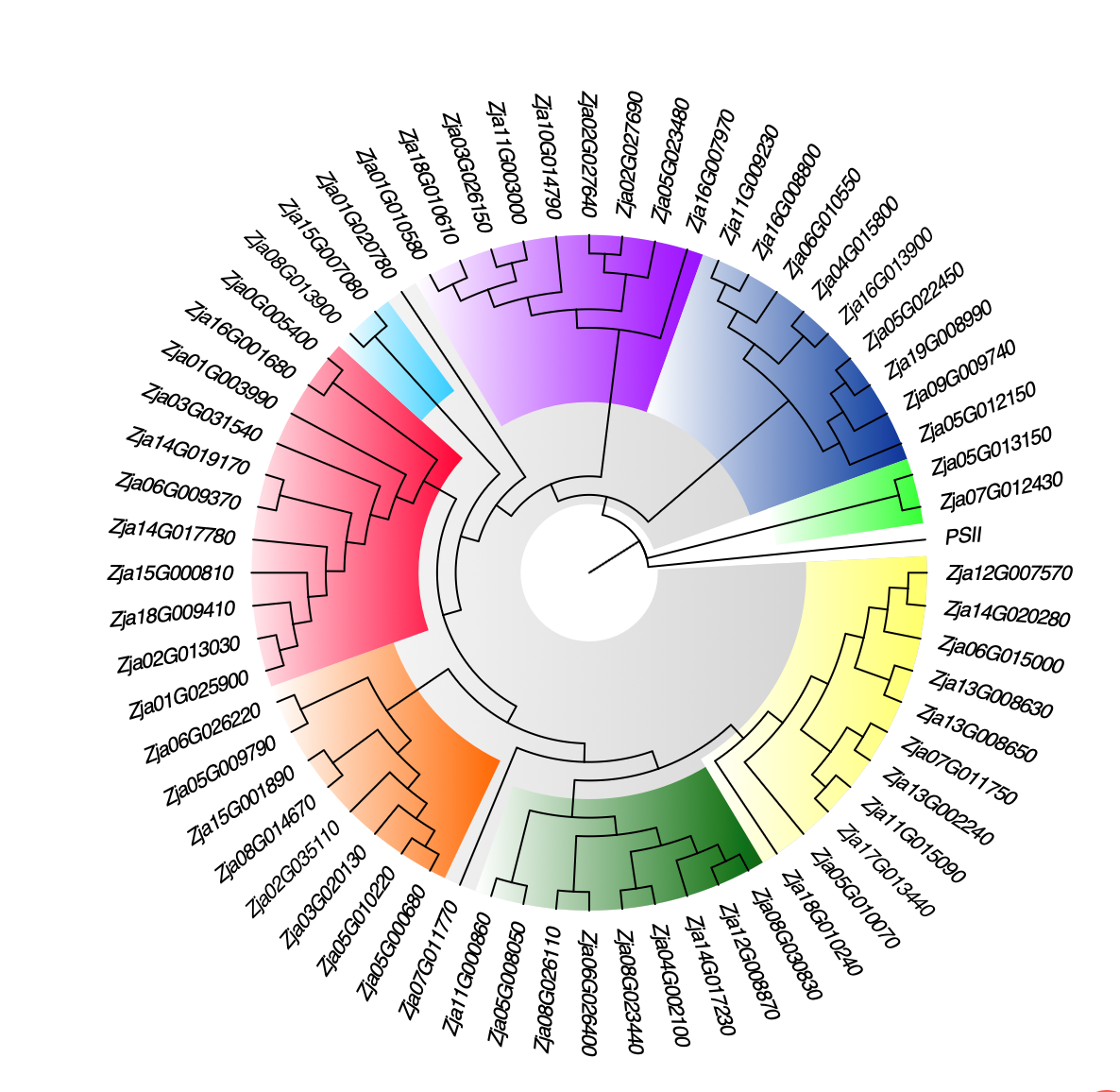

**Fig.S6.** Drought-resistant candidate genes in zoysiagrass

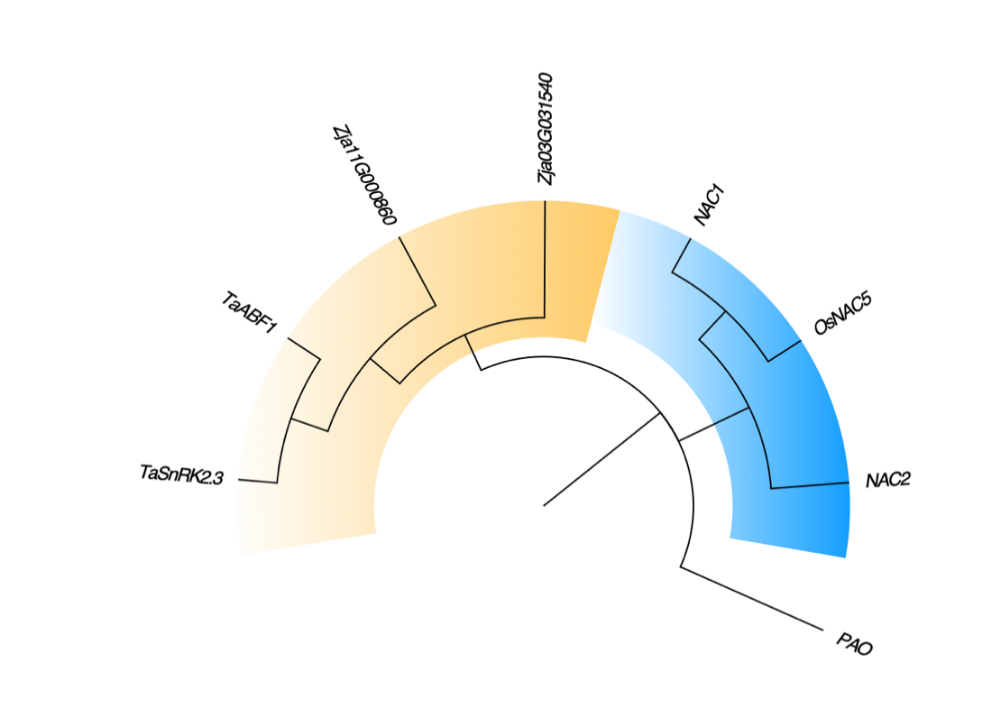

**Fig.S7.** Evolutionary tree of homologous genes. E-value < 1e^-5^.

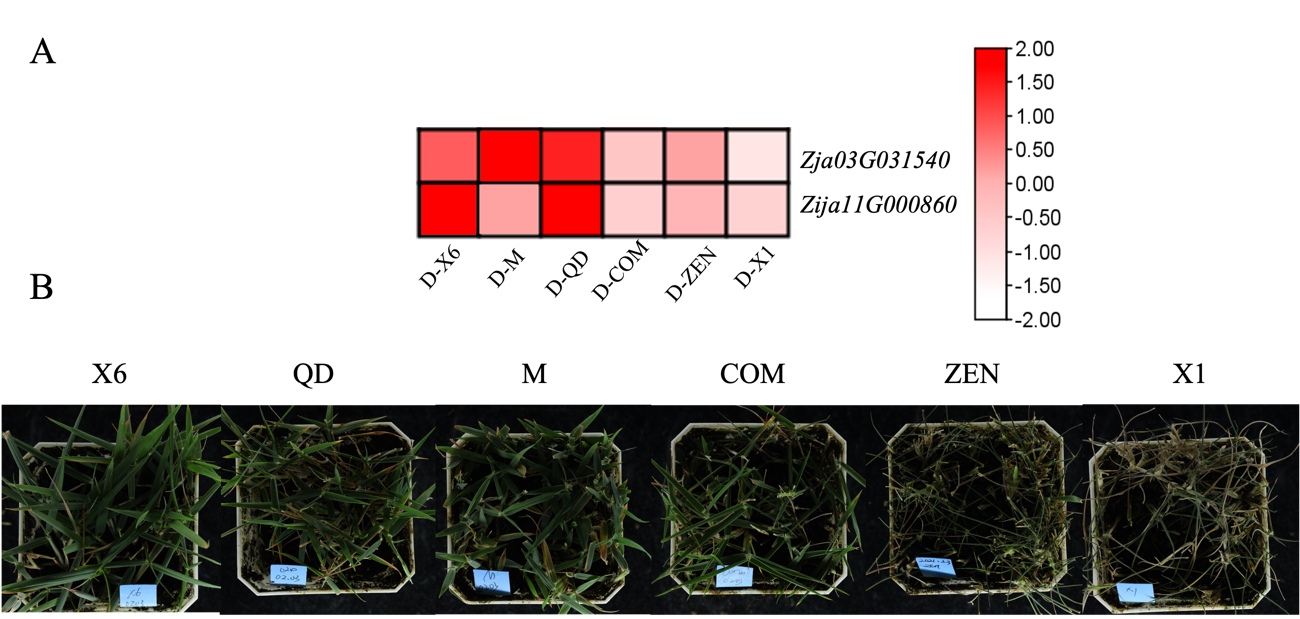

**Fig.S8** The drought response of zoyssiagrass. (A) Expression of candidate drought-resistant genes in zoysiagrass. (B) Phenotypic images of zoysiagrass on the 20th day under drought stress.

Tab.1 Data and analysis of ATAC-seq in zoysiagrass.

| Sample name | CK dataset | CK peak | D dataset | D peak |
| --- | --- | --- | --- | --- |
| COM | 9.45 G | 16724 | 6.24 G | 14630 |
| M | 5.21 G | 15581 | 4.21 G | 14951 |
| QD | 9.86 G | 22790 | 6.18 G | 3134 |
| X1 | 4.82 G | 12618 | 4.29G | 7684 |
| X4 | 4.21 G | 19412 | 4.65G | 13347 |
| ZEN | 6.03 G | 30605 | 9.21 G | 19484 |

Tab.2 Drought-related genes and its information of RT-qPCR in zoysiagrass.

| **Serial Number** | **Gene ID** | **Primer qPCR-F**  **（5’ - 3’）** | **Primer qPCR-R**  **（5’ - 3’）** | **Relative Expression Level** | **Variance** |
| --- | --- | --- | --- | --- | --- |
| D01 | Zja01G003990 | CAAGCCTTACAACCCGGGAT | CAGCTTGAGCAGGATGTCCA | 3.8816 | 0.0461 |
| D02 | Zja01G010580 | GGTGGTGTCGTTTCGGTACT | CAGGCTACATGGTCGGAGTC | 2.0288 | 0.5793 |
| D03 | Zja01G020780 | GGTGAGAAGCCAGTTCGTGA | AGCAGAGAGAGCACTGGAGA | 3.5109 | 1.0031 |
| D04 | Zja01G025900 | ACCGCCCACAGTTCTACATG | GTGTGCCTGTGCCAAATCAG | 3.3368 | 0.0021 |
| D05 | Zja02G013030 | CAACATTAAGCGCGGGGATG | CCATCTTGACACGGTCACCA | 2.0836 | 1.3942 |
| D06 | Zja02G018680 | CTCTCTCCTCGACCCACAGA | TCAAATTCGCGCAAACAGCA | 1.6369 | 2.7372 |
| D07 | Zja02G027640 | TATCCTGCGTCGACATCAGC | AAATCAGCTGGACAGGAGCC | 0.4249 | 2.3189 |
| D08 | Zja02G027690 | TATCCTGCGTCGACATCAGC | AAATCAGCTGGACAGGAGCC | 0.5417 | 0.0079 |
| D09 | Zja02G035110 | TCAAATGGCGGCAACAAGTG | ACCTTCTGCATGAGGAAGCC | 2.665 | 0.0251 |
| D10 | Zja03G017960 | CAACAAGACCAGGTGCGTTG | ATGACTTCTTCAGGGCTGCC | 0.6952 | 1.3149 |
| D11 | Zja03G020130 | TCTGGAGGCGTGATGGAGTA | AATGTCCCCCTGCTGATTGG | 7.6413 | 0.0156 |
| D12 | Zja03G026150 | TCCATCCACCAGATCCCCAT | CAGTTCGACGTCCACCTTGA | 5.2667 | 0.1423 |
| D13 | Zja03G031540 | AAAAGGTGCAGTGAACCCCA | AAGCTTCGCAACTCTCCGAA | 0.2148 | 0.4267 |
| D14 | Zja04G002100 | CGGCAGGACATGTCTCTCTC | ATGGGTAAACGCTCACCTGG | 10.7307 | 0.0037 |
| D15 | Zja04G015800 | CGTCATCACCGCCATTCCTT | CCATCTGGAGAACCGAAGCA | 1.4109 | 0.6850 |
| D16 | Zja04G032370 | CCGGCACAAAAGTCATCGAC | TCACCTCGTTCCCCATCTCT | 0.9368 | 0.0531 |
| D17 | Zja05G000680 | CTGTGCCCAGTAGCATCACA | GGACGATGAGGAGATGCAGG | 5.0918 | 1.3206 |
| D18 | Zja05G008050 | CGTTGGCAGACCTTGTTGTG | CCGCTGCCCATCATAAAACG | 4.929 | 0.6838 |
| D19 | Zja05G009790 | TTTGCATGGCACCAATGTGG | GTAGCACCAATGGACCAGCT | 5.9464 | 0.2855 |
| D20 | Zja05G010070 | TCTGCCAAACATGCCTTTGC | AAGGATGTGCGGCTCAATCA | 1.3165 | 0.4657 |
| D21 | Zja05G010220 | GCGACCACCATTTCCTCTCT | GCTTCCTCCACAGTGCTCAT | 1.3318 | 0.1042 |
| D22 | Zja05G012150 | TCGATGATACGACTGCTGCC | GCCAGCCTTATCATCGTCGA | 4.685 | 0.0663 |
| D23 | Zja05G012980 | CCCAAGACTGCCGAGAACTT | TTTCGTGAACGCTGGTCAGA | 0.4545 | 1.9386 |
| D24 | Zja05G013150 | CTGACCGGGAAGACCATCAC | GCCATAGAGTTACCGAGCCC | 7.6393 | 0.6225 |
| D25 | Zja05G022450 | TCTGGGAGAGCTTTGGCAAG | CCTTGTCACCAAGCTGTTGC | 2.5741 | 0.6783 |
| D26 | Zja05G023480 | GCAAGCGAGATAAAGGCTGC | TGCCCATCACCAAAACCCTT | 3.4511 | 0.0059 |
| D27 | Zja06G009370 | CGGAGCCTCTTAGCCTAAGC | AGGACCAGCTCCCATCAGAT | 5.5151 | 0.0136 |
| D28 | Zja06G010550 | TAGATTCGTCGCGTTACCCG | GACGACAAAACTGGTGTGGC | 13.2061 | 0.0066 |
| D29 | Zja06G015000 | TCCTCTTCCTCCTCGACCTG | AGGTGTTCACTACTGGCGTG | 1.743 | 0.3078 |
| D30 | Zja06G026220 | CGTCGTGTTGAACCGTTGAC | TCTGATCGCCTTCTCCAACG | 1.4613 | 1.8192 |
| D31 | Zja06G026400 | AGCCGTGAGAGGAAGATTGC | GGACACCGATCATTGCAAGC | 4.0855 | 0.6063 |
| D32 | Zja07G011750 | GTCCAGACCGCTAGTTTGCT | ACAGCTGCATTATGGTGCCT | 25.8072 | 0.0205 |
| D33 | Zja07G011770 | CACAGGCAGATGACCGAAGT | CAAGCTGGAGTAAGCCACGA | 2.3341 | 0.5923 |
| D34 | Zja07G012430 | CGGGTTCGTCAAGATGCAGA | TCCAATTCCATCCGAGCACA | 3.403 | 0.7228 |
| D35 | Zja08G005310 | AAAGGGTTCCTCGCGCTG | GCTCCCTCTTCTGCAATGGT | 3.9652 | 0.0257 |
| D36 | Zja08G013900 | ATCAGGGGGAAGAAGGTGGA | CCGTTCGCACAGCAATTGAT | 1.9084 | 0.8339 |
| D37 | Zja08G014670 | TTCGGTGACAGCAGGAACTC | CTGTCCCCGGAATCCAGTTC | 1.8575 | 1.4550 |
| D38 | Zja08G023440 | TTCTCTCCTTCCCGTGTCGA | TCAAGCCACCGTGACCAATT | 1.9243 | 0.0442 |
| D39 | Zja08G026110 | TATGGGTCTGCTGCTATGCG | TGCTTACTGACACAGGCAGG | 2.4587 | 0.1527 |
| D40 | Zja08G030830 | GGCTCTAGTCAAGGCTTGCA | GCACTTCCCTTTGAGAGCCT | 2.2734 | 1.1114 |
| D41 | Zja09G009740 | GCCTTGTGTCCGTTAGGTCA | GGCGAGAATCCCACATCCAA | 2.8473 | 0.0172 |
| D42 | Zja0G005400 | GATCTCCTGCATCGGATGGG | GTCGAAGAAGCCCTGGTTGA | 1.1837 | 0.0404 |
| D43 | Zja10G014790 | AGCATAGCCTTGGGAACACC | TGCCTTTAGAGACAGCAGCC | 12.4944 | 0.4863 |
| D44 | Zja11G000860 | GGCCATGTAAGGATCTCGGG | ACGACCACGCAATAAGAGCA | 0.6267 | 0.8027 |
| D45 | Zja11G003000 | CTAGAGGTTGTGCTCGGAGC | GCTTCATGCCCAAAATCGCT | 4.4557 | 1.1426 |
| D46 | Zja11G009230 | GGCGAGTAGGAGTCAGTTCG | ATGTCACTAGTGCCACCTGC | 1.875 | 0.0035 |
| D47 | Zja11G015090 | TTCCCAGATCCCACGTTGTG | GCAATGACAACCAGATCCGC | 3.1672 | 0.0184 |
| D48 | Zja12G007570 | CGAGGAGTCCATACATGCCC | CGCCGGTCGAATCTACATCA | 4.2502 | 0.1798 |
| D49 | Zja12G008870 | AAGCAACTCTCATGGCCCTC | ACCATCAAGAAGGCCTCGTG | 3.4822 | 0.4441 |
| D50 | Zja13G002240 | CTCCCTGGCTGAAGAAGTGG | CAGTCTTCATGTCATGCGCG | 1.5476 | 0.3657 |
| D51 | Zja13G008630 | CCTCCCCTATTGTTCGAGCC | GGGGCGGTAGGATCAAGATG | 1.8485 | 6.6360 |
| D52 | Zja13G008650 | GGTGGGTTCATCATGGTCGT | ACTTGTCAACGCATGCAACC | 4.149 | 0.4327 |
| D53 | Zja14G017230 | TCACCTCTCTCTTCCTCCCG | CCACGCCTTCCAGTATCTCC | 2.9772 | 0.0029 |
| D54 | Zja14G017780 | CAAGGTGGTTGCCATTGGTG | TTCTCCATCATGGCGACGAG | 4.2575 | 3.7851 |
| D55 | Zja14G019170 | GTGTTGGGCCACGACTTTTC | GAGTAGCGATCACTCCTGGC | 0.1119 | 0.0285 |
| D56 | Zja14G020280 | AGCCCTATTTCTGGTGTGGC | GCTGAAACACACCCTGCAAG | 7.9081 | 0.0005 |
| D57 | Zja15G000810 | CTCTCTTCGCCAAGCTCCTC | GCACCACCTCCACCTACATT | 3.491 | 0.1224 |
| D58 | Zja15G001890 | CCGCTCTACCTTGCCGTATT | AGACAACCTTCAGCTCCTGC | 4.245 | 0.7082 |
| D59 | Zja15G007080 | GCACCCATCTCCTATCGACG | CGGGGAGCATGATTACCCAG | 4.4393 | 1.1596 |
| D60 | Zja16G001680 | AGCGAGGACTTTGCAGAGAC | CTCGTATCCCACAGCTACGC | 1.8674 | 0.0069 |
| D61 | Zja16G007970 | GCAATAAGCTTGGGCGAGTG | TGTTGTGTCAAGCCAACCCT | 2.9485 | 0.0857 |
| D62 | Zja16G008800 | GCTGAGAAGAGAGGGAAGCG | AAACCGAGAACCTTGCCTCC | 20.0237 | 0.3274 |
| D63 | Zja16G013900 | TGCTCAAGCCTCAACTCCTG | TCCCCTGGAGAGGTGCTATC | 2.6421 | 0.3722 |
| D64 | Zja17G013440 | TCATCATGGACGTCGTCACC | CTTGGTCTGCATGACGAGGT | 1.8165 | 0.1543 |
| D65 | Zja18G009390 | CGTGCTGCATTCTTCATCCG | GATCGGCACCTGAGAGGATG | 0.7329 | 0.1372 |
| D66 | Zja18G009410 | TCCGGCTCTTCTGGTCAAAC | CTGTGTGGAAGAGGTTCGCT | 1.7581 | 0.0097 |
| D67 | Zja18G010240 | CCAAGAAGAGGAGGGGGAGG | TCCACCCAAAACAGCCTACC | 2.1873 | 0.0014 |
| D68 | Zja18G010610 | CTGGCAAGCCTACATCCACA | GCCTCGTCCATAAGAGCTCC | 0.7347 | 0.8914 |
| D69 | Zja19G008990 | CAGACTGCATGTGTGTCCCT | ATCCGGTGCCTTAGAGTCCT | 0. 0472 | 0.1034 |
| D70 | Zja20G005590 | ATGGGTGCTTCCTGGGAATG | CAGATTCCACGTGGAGGCTT | 2.6259 | 0.0577 |
